# Evaluation of five cell-free DNA isolation kits for plasma

**DOI:** 10.1101/715821

**Authors:** Zhongzhen Liu, Xi Yang, Haixiao Chen, Sujun Zhu, Juan Zeng, Fang Chen, Wen-Jing Wang

**Affiliations:** BGI-Shenzhen, Shenzhen 518083, China; China National GeneBank, BGI-Shenzhen, Shenzhen, China; Obstetrics Department, Shenzhen Maternity and Child Healthcare Hospital, Shenzhen, Guangdong Province, China

**Keywords:** cell-free DNA, isolation kits, recovery efficiency, liquid biopsy

## Abstract

Cell-free DNA (cfDNA) has been widely used in prenatal test and cancer diagnosis nowadays. The cost- and time-effective isolation kits are needed especially in large-scale clinical application. Here, we compared three domestic kits: VAHTS Serum/Plasma Circulating DNA kit (VZ), MagPure Gel Pure DNA mini kit (MG) and Serum/Plasma Circulating DNA Kit (TG), together with QIAamp Circulating Nucleic Acid Kit (QC) and QIAamp DNA Blood Mini Kit (QD) in cfDNA isolation. cfDNA was isolated from the pooled samples with spike-in fragments, qPCR was conducted to quantify the spike-in fragments recovery. The results indicated that all of the five kits could isolate cfDNA with different efficiency. The VZ kit had an efficiency as high as 90 percent, which is comparable to QC kit. The libraries were constructed using the isolated cfDNAs, quantified by Qubit and analyzed by 2100 bioanalyzer. Both showed the libraries were qualified. Finally, cffDNAs were detected by qPCR targeting SRY gene using libraries from pregnant women bearing male fetuses. All five kits could isolate cffDNAs that could be detected by qPCR. Our results provided more choices in wide-scale clinical application of cfDNA-based non-invasive genetic tests.

## Introduction

Since the discovery of the circulating tumor DNA from cancer patients serum and plasma in 1996 and fetal DNA from maternal serum and plasma in 1997, the circulating cell-free DNA (cfDNA) has gotten widely attention [1–3]. The cfDNAs originate from the apoptosis, necrosis or active release of tumor cells in cancer patients, fetal trophoblastic cells in pregnant women, or donor cells after transplantation [4]. Due to the safety and convenience, cfDNAs have been widely used as biomarkers in non-invasive fetal screening (NIPS), organ transplant graft rejection [5], trauma [6], sepsis [7], myocardial infraction [8].

In general, there are two common ways through which the analysis of cfDNAs are conducted. One way is quantified PCR, including qPCR or digital PCR, which can analyze the target genes with site mutations or even structural variant. The limitation is the mutations or SVs should have been proven to potentially involved in specific cancer or birth defects [3, 9]. The second way is next generation sequencing, which is competent to detect all of the potential mutations or SVs of the cell-free tumor DNA (ctDNA) or cell-free fetal DNA (cffDNA) from very early stagy, and give valuable information for diagnosis and treatment [10, 11].

One of the most important steps of cfDNA analysis is extraction of cfDNAs from the liquid biopsies. It is known that the concentration of cfDNA in plasma is usually very low, ranging from several to tens of nanogram per milliliter [12]. The variety of the concentration is large according to the progression of disease or pregnant terms. Moreover, the ‘valuable cfDNA’, which means the ctDNA from the tumors or cffDNA from the fetuses only accounts for small proportion of the total cfDNA, especially in early stages at which the diagnosis is more valuable. Given the samples are limited, the efficient cfDNA extraction kits are needed to be developed. Nowadays the most popular kits are QIAamp Circulating Nucleid Acid kits, which can give a stable and high recovery, while the higher cost of the kits limits the massive application. The traditional Triton/Heat/Phenol (THP) method is cost-effective, but costs much time, and is difficult to conducted automatically.

A series of reports compared the homemade or commercially available cfDNA extraction kits [13–18], which gave valuable guidelines to select the suitable kits for different samples. Still there are limitations in these studies. Most of the kits are less suitable in industrial application due to the time- and money-consumption. In the present study we directly compared three domestic cfDNA extraction kits with QIAamp Circulating Nucleid Acid kits using real-time PCR and Agilent 2100 bioanalyzer. We found that recovery efficiency of cfDNAs by at least one kit was comparable to QIAamp while more cost-effective. Our results gave an additional choice in cfDNA isolation.

## Materials and methods

### Plasma sample collection and pretreatment

Five mL peripheral blood were taken using EDTA anticoagulant-coated tubes from pregnant women in the second trimester. All blood samples were centrifuged at low speed (3000 rpm) for 5 min at 4□ within three hours after collection. The supernatant was centrifuged at high speed (1,4000 rpm) for 15 min at 4□. The genders of the fetuses were indicated by ultrasound and confirmed by PCR using plasma. The blood samples were fell into three groups: for group I, 1 mL plasma from six pregnant women bearing female fetuses were pooled together and divided into 15 aliquots and processed by five kits. For group II, 1 mL plasma from the identical six pregnant women were pooled together and 15,000,000 copies of spike-in-162, 340, 500 and SRY were added, and then divided into 15 aliquots and processed by five kits. For group III, 2.5 mL plasma from three pregnant women bearing male fetus was processed by these five kits separately. Written consent forms were obtained from all women, and the study was approved by the BGI Institutional Review Board (BGI-IRB17166).

### Spike-in fragments preparation

The spike-in fragments were constructed by insertion the SPUD double strand DNA (5’-AACTTGGCTTTAATGGACCTCCAATTTTGAGTGTGCACAAGCTATGGAACACCACGTAAGACATAAAACGGCCACATATGGTGCCATGTAAGGATGAATGT-3’) into pCE2-TA/Blunt-Zero vector using 5 min TA/Blunt-Zero Cloning Kit (vazyme, C601). The amplicons with different lengths (162, 340, 500bp) from pCE2-TA/Blunt-Zero vector were amplified using the primer pairs: spike-in-162 Fw & Re, spike-in-340 Fw & Re or spike-in-500 Fw & Re. SRY segments was amplified with primer pairs spike-in-SRY Fw & Re using male genomic DNA as template. After GEL-purification and quantification, 15,000,000 copies of each fragment were added in aliquots in group II. All the primers were listed in table 1.

**Table 1.**
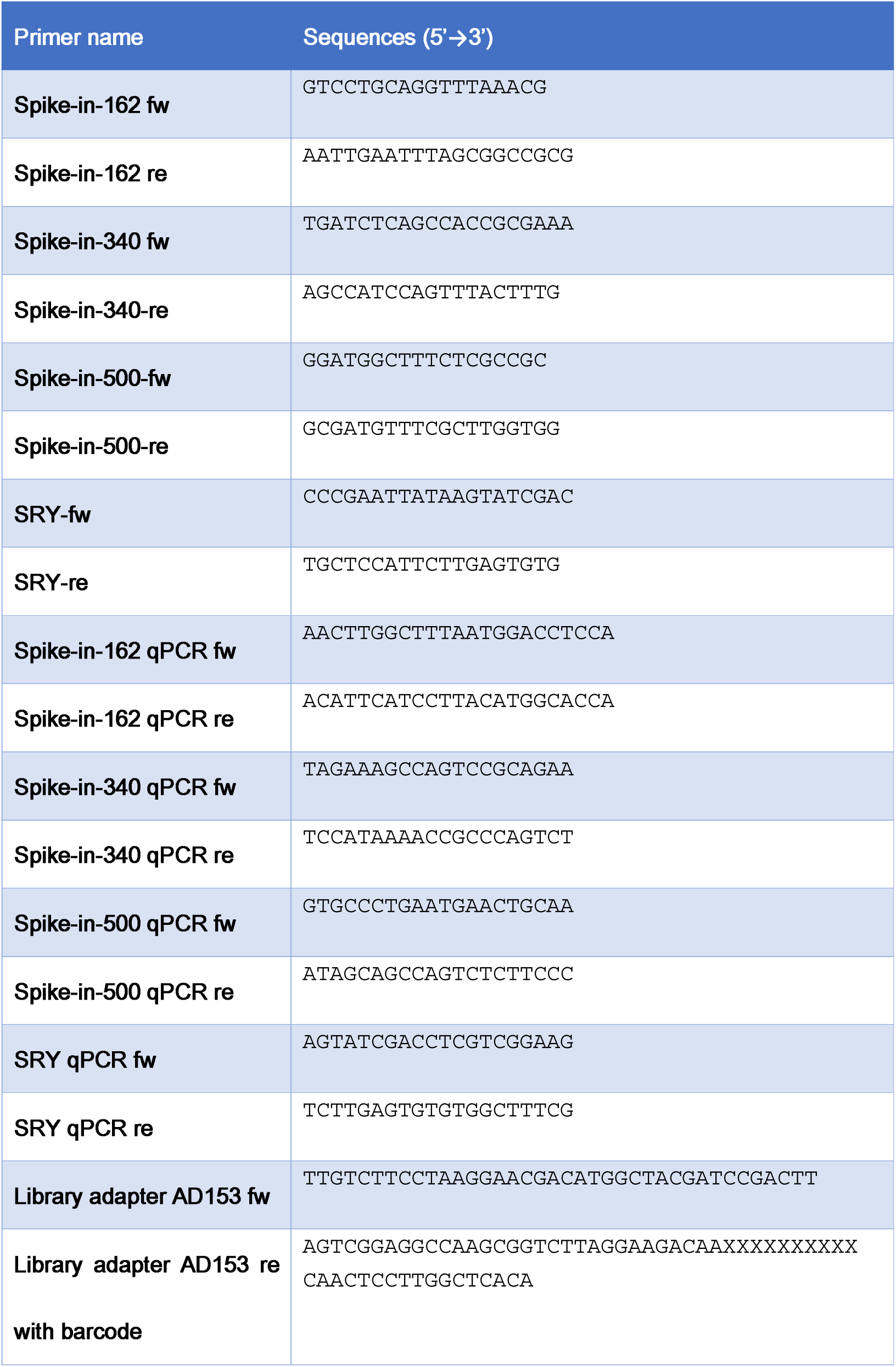
primers used in this study

### DNA extraction

The cfDNA was extracted from 200 uL plasma using VAHTS Serum/Plasma Circulating DNA kit (Vazyme, Cat. No.: N902-01-BOX2, VZ for short), MagPure Gel Pure DNA mini kit (Magen, Cat. No.: MD5001-02, MG for short), Serum/Plasma Circulating DNA Kit (TIANGEN, Cat. No.: DP339, TG for short), or from 1 mL plasma using QIAamp Circulating Nucleic Acid Kit (QIAGEN, Cat. No.: 55114, QC for short) following the manufacturer’s guide. We also used QIAamp DNA Blood Mini Kit (QIAGEN, Cat. No.: 51104, QD for short), which was not designed to extract cfDNA. The cfDNA was eluted by 200 uL TE buffer for QC and 40 uL for the rest.

### Quantitative PCR

qPCR was conducted on StepOnePlus™ Real-Time PCR System (Applied Biosystem, 4376600) using SYBR Premix Ex Taq (Tli RNaseH Plus), ROX plus (Takara, Cat. No.: RR420LR). The primers for qPCR were listed in table 1.

### cfDNA library construction

The extracted cfDNAs were processed to library using MGIEasy Cell-free DNA Library Prep kit (MGI, cat. No.: AA00226). The Ad153 Fw & Re adapters listed in table 1 were ligated to the amplicons.

### Qubit DNA quantification

The library was qualified using Qubit dsDNA HS Assay Kit (Invitrogen, cat. No.: Q32851). 2 uL of each library was loaded and the concentration was detected by Qubit 3.0 Fluorometer (Invitrogen).

### 2100 Bioanalyzer assay

The libraries were loaded on DNA 1000 series II chip to undergo automated electrophoresis using Agilent 2100 Bioanalyzer (Agilent) for sizing and qualification of the library.

## Results

### The cfDNA yield was comparable between VZ and QC, with the efficiency more than 90%

To compare the cfDNA yield directly, we pooled six samples together and divided into 15 tubes, 200 uL each for the VZ, MG, TG, QD and 1 mL each for QC, as the lowest volume as the kit requires. We extracted cfDNA with five kits, each kit had three repeats. Due to the addition of carrier nucleic acids, which could increase the recovery according to the product manual of TG, QC and QD, the concentration of cfDNA in elution could not be detected directly. To overpass this problem, we evaluated the cfDNA yield by using exogenous spike-in fragments. We added a mixture of spike-in-162, 340, 500, spike-in-SRY (162 bp in length), 1,500,000 copies for each segment, to 200 uL plasma or 7,500,000 to 1mL plasma for QC. We detected the number of copies of each segment in elution buffer using qPCR, and calculated the recovery efficiency (RE) by using the equation RE = copy number eluted / 1,500,000 (7,500,000 for QC). As shown in figure 1, generally, the RE of QC was highest for all four segments (100.64%, 111.45%, 92.41%, 68.81%). The VZ kit displayed similar yield (96.54%, 95.03%, 87.93%, 73.49%). The RE of MG kit was lower, while the TG kit was the lowest. The QD kit, which is not designed to isolate cfDNA, also generated weak signals.

**Figure 1.**
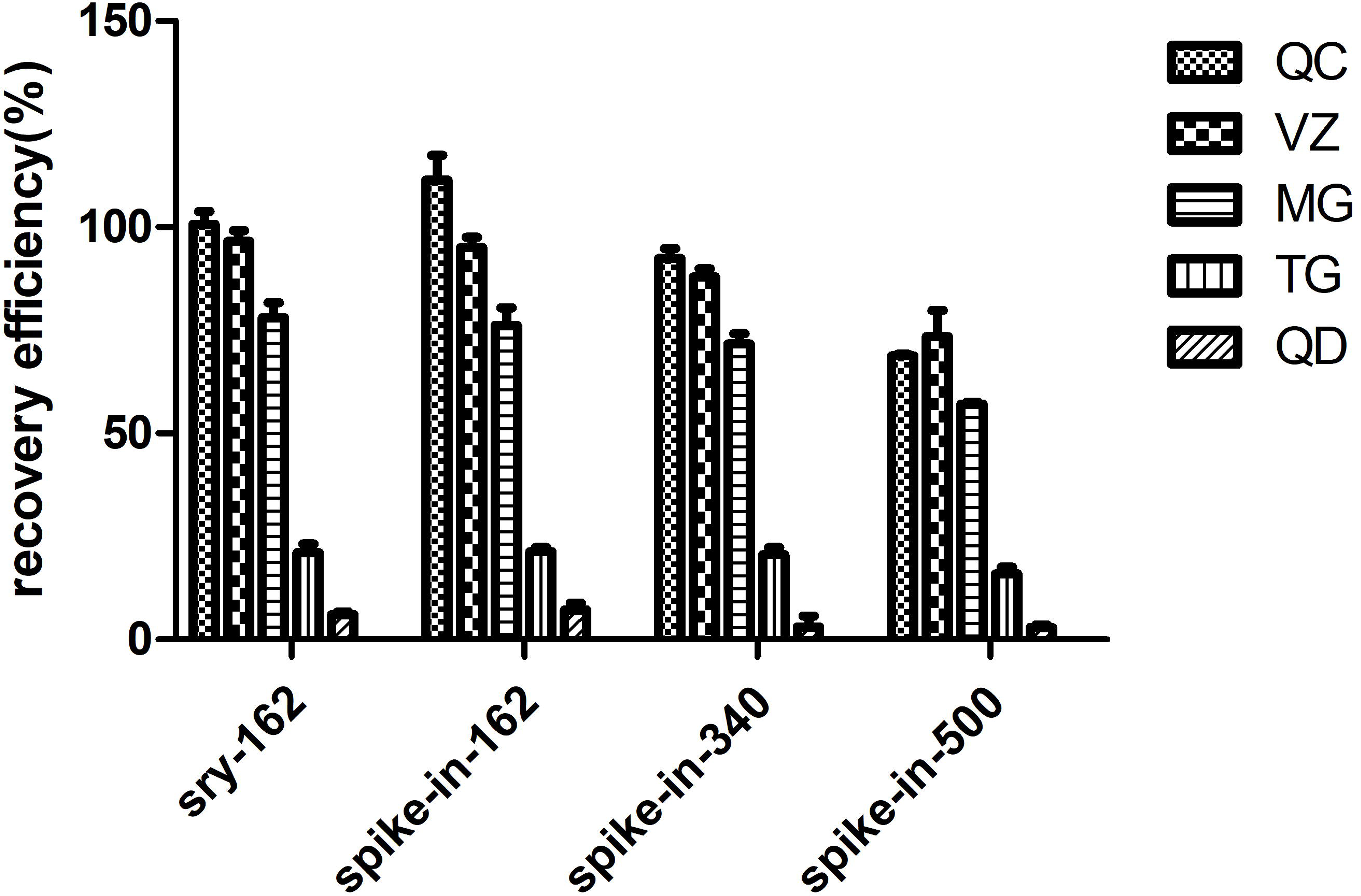
Comparison of five cfDNA isolation kits by using the percentage of spike-in recovery generated by VZ, MG, TG, QC, QD. Four spike-in fragments with indicated lengths were added in the pooled plasma. Three replicates were conducted for each kit. qPCR was conducted to calculate the copy numbers using standard curve method. Note that for some fragments the efficiency was higher than 100 percent, probably due to the systematic errors of qPCR.

There was a relationship between RE and size of segments. The yields of spike-in-162 and spike-in-SRY, both of which are 162bp, was the highest for all of the four kits. On the other hand, spike-in-500 had the lowest RE. It has been proven that 162 bp fragments account for the majority of circulating DNA, thus for QC and VZ kits, nearly 90% of cfDNA could be isolated.

### The isolated cfDNA was qualified for library construction

Next, we evaluated whether the cfDNAs isolated through these five kits could be used in library construction for NIPT. We extracted cfDNAs from the plasma in the group I using different kits and kept 20 uL elution to conduct library conduction. We eluted the libraries in 20 uL elution buffer, and evaluated the concentration of the libraries.

As shown in figure 2, cfDNAs from all five kits were qualified to construct library (figure 2). As in spike-in experiments, the concentration of libraries from VZ and QC has the highest concentration, nearly 30 ng/uL. MG and TG kits can reach to more than 10 ng/uL.

**Figure 2.**
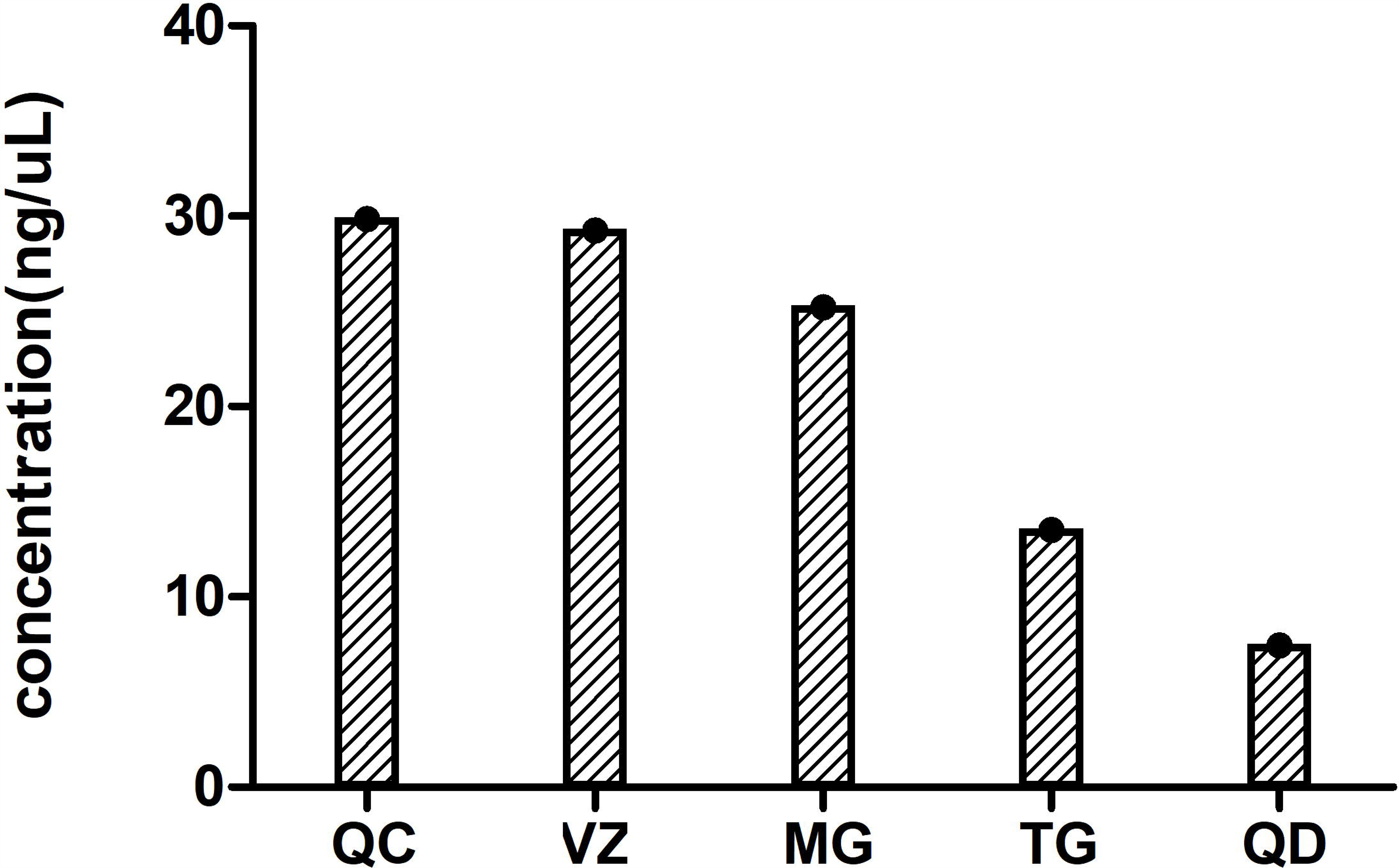
The concentration of libraries constructed from cfDNAs isolated by five kits from pooled plasma. Three replicates were conducted for each kit. And 20 uL elution was used to construct libraries.

We also analyzed the molecular weight and concentration of libraries using 2100 bioanalyzer. As expected, there was a main peak at around 250 and a minor peak at around 430 (figure 3). The tendency was similar with spike-in experiment.

**Figure 3.**
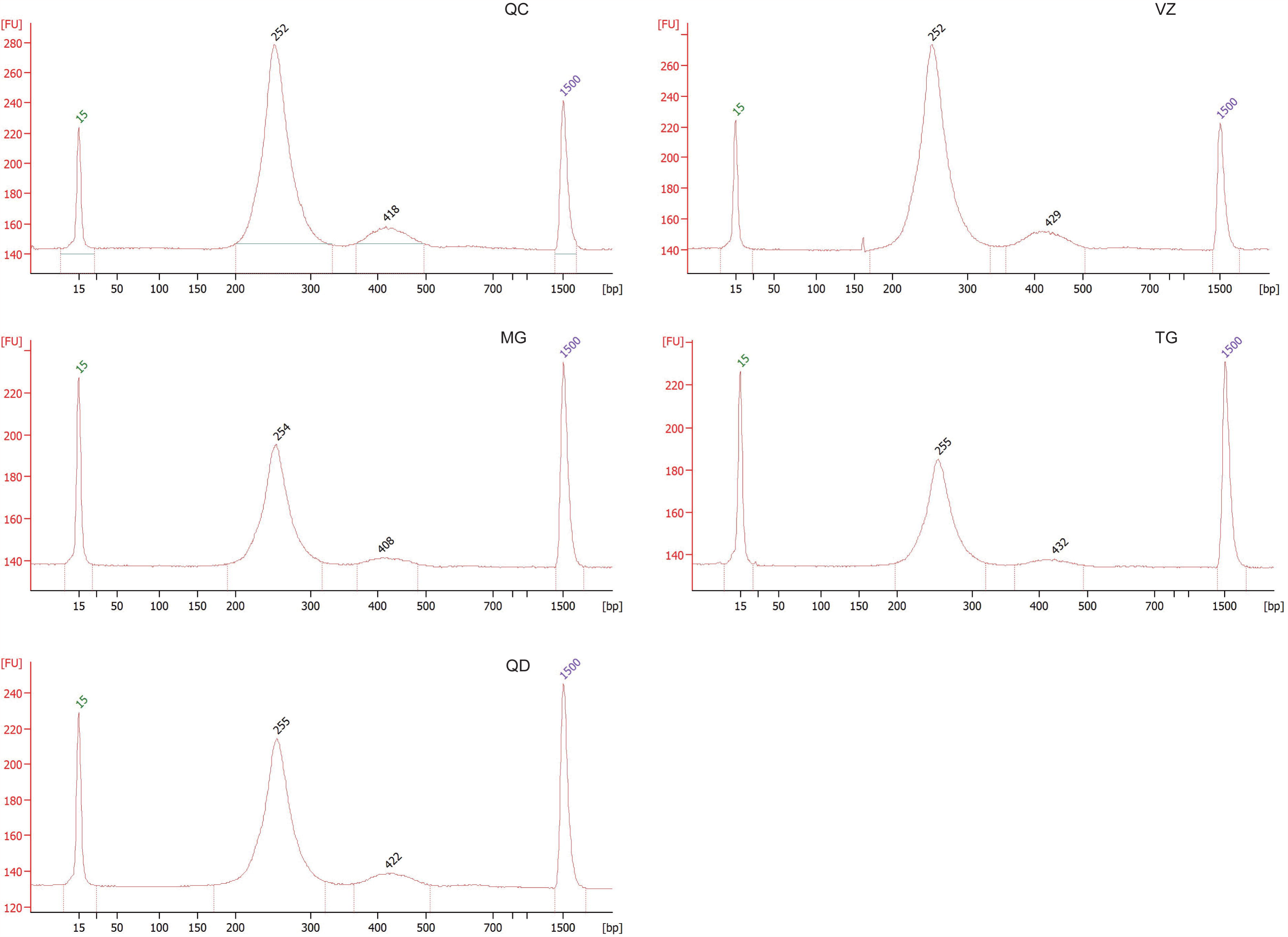
The 2100 bioanalyzer results of libraries constructed from cfDNAs isolated by five kits from pooled plasma. One library of each kit was loaded.

### All of the three kits were efficient to detect cell free fetal DNA

Due to scarcity of cffDNA in mother’s plasma, the efficient recovery of cffDNA calls for high efficiency of the kits and no bias between cffDNA and mother’s cfDNA. We evaluated the cffDNA isolation efficiency by detecting copy number of Y chromosomal segments isolated from the plasma of mothers bearing male fetuses. We isolated cfDNA from plasma in group III using these five kits separately and used 20 uL elution to construct libraries. Then we conducted qPCR using the primer pair identifying SRY fragments and the libraries as templates. Our results indicated that the cfDNAs from all five kits could give signals of SRY fragments (figure 4). We can also calculate the copy numbers of SRY segments in the libraries. We found for all three plasma, the SRY copy numbers from QC and VZ libraries were more than 5000 copies. The copy numbers from MG and TG were smaller, but still enough to detect the fetal genome.

**Figure 4.**
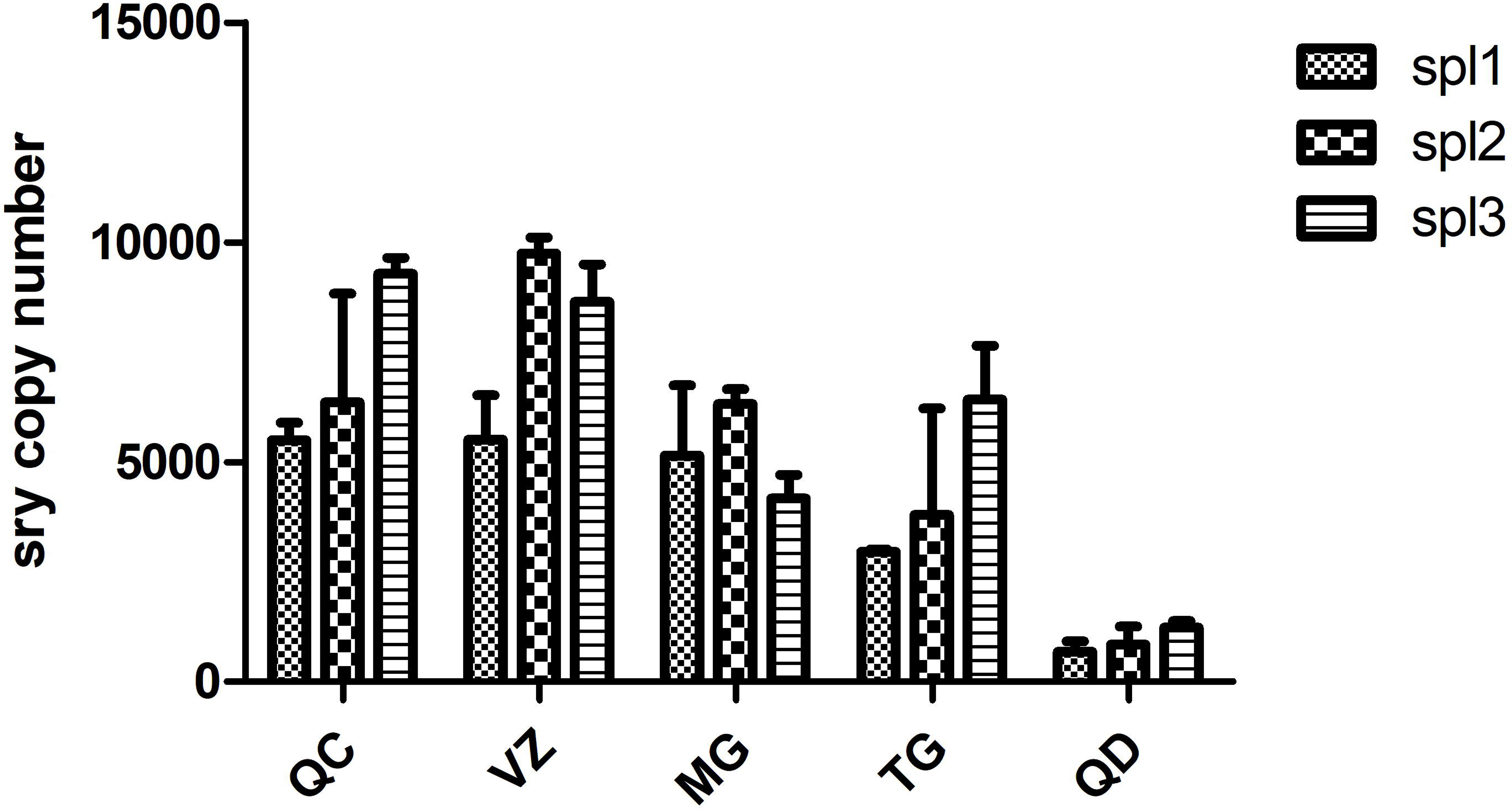
The copy numbers of SRY in 1 uL libraries constructed from cfDNA isolated by five kits from three pregnant women bearing male fetuses.

## Discussion

In this study, we evaluated five isolation kits of cfDNA extraction from plasma of pregnant women. We compared the direct recovery efficiency of these five kits by investigation of exogenous spike-in fragments using qPCR and found all five kits could extract cfDNAs with different efficiency, while the recovery of QC and VZ could reach up to 90%. We also evaluated the libraries constructed from the cfDNAs extracted by these five kits, and all were qualified for library construction.

cfDNA analysis has been proven a powerful tool in cancer diagnosis and NIPS. As the initial step of cfDNA analysis, cfDNA extraction is the most important, because the purification and recovery efficiency determines the following analysis steps and finally the signal quantity. Nowadays there were a series of studies investigating the efficiency of several kits, including QIAamp circulating nucleic acid kit (QIAgen, Valencia, CA, USA), MagNA Pure Compact (MPC) Nucleic Acid Isolation Kit I (Roche Diagnostics, Penzberg, Germany) etc., which are popular worldwide. As the large-scale clinical application of cfDNA analysis developed recently, the lower-cost as well as efficiency-comparable kits are in need, especially in the less developed countries, like China. Thus, in this study, we focused on three domestic DNA extraction kits, VZ, MG and TG, which are less expensive.

Another factor that influences large-scale application is the automation. At present there are two technologies employed in commercial cfDNA purification kits: spin column-based and magnetic beads-based approach. Compared to column, the magnetic beads-based approach could be conducted in 96-well plates automatically and has been applied in large-scale cfDNA extraction. Indeed, the VZ and MG kits has already been used in automatic extraction platform and performed stably.

Notably, the recovery efficiency of spike-in calculated by qPCR was higher than 100%, probably due to the systematic error of qPCR, which is not easily avoided. The more exact method was digital PCR, which we are trying to establish.

It has been proven that there are a dominant peak at approximate 162 bp and a minor peak at around 340 bp [19]. We did not run 2100 after cfDNA extraction directly due to addition of carrier nucleic acids according to the manual of TG, QC and QD. Alternatively, we run 2100 after library construction, when the 90bp adapters were added to the cfDNAs. There should be a major peak at ~250bp and a minor peak at ~430bp. Our 2100 bioanalyzer results were in accordance with this conclusion. It is reasonable to use 162bp-, 340bp-, 500bp-spike-in fragments, which represent the main parts of the cfDNAs, in our experiments. For QC and VZ, the recovery efficiency of 162bp and 340bp fragments was more than 90%, so we can give a bold prediction that these two kits have a RE of at least 90 percent.

As NIPS is widely used in prenatal screening, the cost- and time-efficient cfDNA extraction methods are valuable in widescale application. Our results may provide more choices in a clinical setting.

## Acknowledgement

We are grateful to the participants who donated their samples to our project. This project is supported by the National Natural Science Foundation of China (No.81300075), the Natural Science Foundation of Guangdong Province (No. 2014A030313795), the Shenzhen Municipal Government of China (No.JCYJ20170412152854656, JCYJ20180703093402288).

